# Accurate, Sensitive, and Precise Multiplexed Proteomics using the Complement Reporter Ion Cluster

**DOI:** 10.1101/205054

**Authors:** Matthew Sonnett, Eyan Yeung, Martin Wühr

**Affiliations:** Department of Molecular Biology and the Lewis-Sigler Institute for Integrative Genomics, Princeton University, Princeton, NJ, USA

## Abstract

Quantitative analysis of proteomes across multiple time points, organelles, and perturbations is essential for understanding both fundamental biology and disease states. The development of isobaric tags (e.g. TMT) have enabled the simultaneous measurement of peptide abundances across several different conditions. These multiplexed approaches are promising in principle because of advantages in throughput and measurement quality. However, in practice existing multiplexing approaches suffer from key limitations. In its simple implementation (TMT-MS2), measurements are distorted by chemical noise leading to poor measurement accuracy. The current state-of-the-art (TMT-MS3) addresses this, but requires specialized quadrupole-iontrap-Orbitrap instrumentation. The complement reporter ion approach (TMTc) produces high accuracy measurements and is compatible with many more instruments, like quadrupole-Orbitraps. However, the required deconvolution of the TMTc cluster leads to poor measurement precision. Here, we introduce TMTc+, which adds the modeling of the MS2- isolation step into the deconvolution algorithm. The resulting measurements are comparable in precision to TMT-MS3/MS2. The improved duty cycle, and lower filtering requirements make TMTc+ more sensitive than TMT-MS3 and comparable with TMT-MS2. At the same time, unlike TMT-MS2, TMTc+ is exquisitely able to distinguish signal from chemical noise even outperforming TMT-MS3. Lastly, we compare TMTc+ to quantitative label-free proteomics of total HeLa lysate and find that TMTc+ quantifies 7.8k versus 3.9k proteins in a 5-plex sample. At the same time the median coefficient of variation improves from 13% to 4%. Thus, TMTc+ advances quantitative proteomics by enabling accurate, sensitive, and precise multiplexed experiments on more commonly used instruments.

## Introduction

Global measurements of protein abundance are essential to understanding biological systems in health and disease. However, proteomics measurements severely lag behind other “omics” approaches such as transcriptional profiling.^1^ Proteomic measurements lack sensitivity, lack throughput, are comparatively expensive, and can produce unreliable quantification.

The majority of modern proteomics relies on two widely employed techniques, label-free proteomics and multiplexed proteomics. In label-free proteomics, samples for individual conditions (e.g. time-points or perturbations) are analyzed one at a time, peptide ion-intensities are mapped to proteins, and the resulting intensities are compared between different experiments. However, due to the complexity of the samples even the fastest instruments cannot fragment and identify all peaks.^2^ Peptide (and thus protein) identification among different conditions is therefore a somewhat stochastic process. Comparing different samples analyzed with label free produces missing data-points which complicate and hinder interpretation.

Furthermore, typically only two-fold or larger changes can be detected as significant.^3^ Despite these disadvantages, the comparative ease of implementation and compatibility with comparatively simple and robust instrumentation have made the label-free approach highly attractive and widely used.

In principle, multiplexed proteomics can address many of the shortcomings of label-free proteomics. Multiplexing multiple conditions into a single mass spectrometer run is accomplished by labeling peptides with isobaric tags (e.g. Tandem Mass Tag (TMT)) that function as barcodes and specify the different conditions (Fig. 1A).^4^ The different conditions are combined and analyzed together, which results in simultaneous ionization, in principle leading to more reliable quantification and the elimination of missing values. Multiplexing increases sample throughput and reduces the need for expensive instrument time. In the MS1 spectrum, intact peptides tagged with different TMT-tags are indistinguishable as each tag is isobaric and thus has the same mass (Fig. 1B). Therefore, the MS1 spectrum complexity does not increase with more channels, enabling the comparison of many (currently up to 11 conditions) in a single experiment.^5^ Tagged peptides are isolated and fragmented in the MS2 spectrum (Fig. 1C). The fragment ions resulting from breakage of the peptide backbone, called b- and y-ions, are used for peptide identification. Additionally, the TMT-tag will break during the fragmentation process and release the reporter-ions; unlike the intact tag, the masses of the resulting reporter-ions are condition specific and encode relative protein abundance between samples (Fig. 1C).^4^

**Figure 1:**
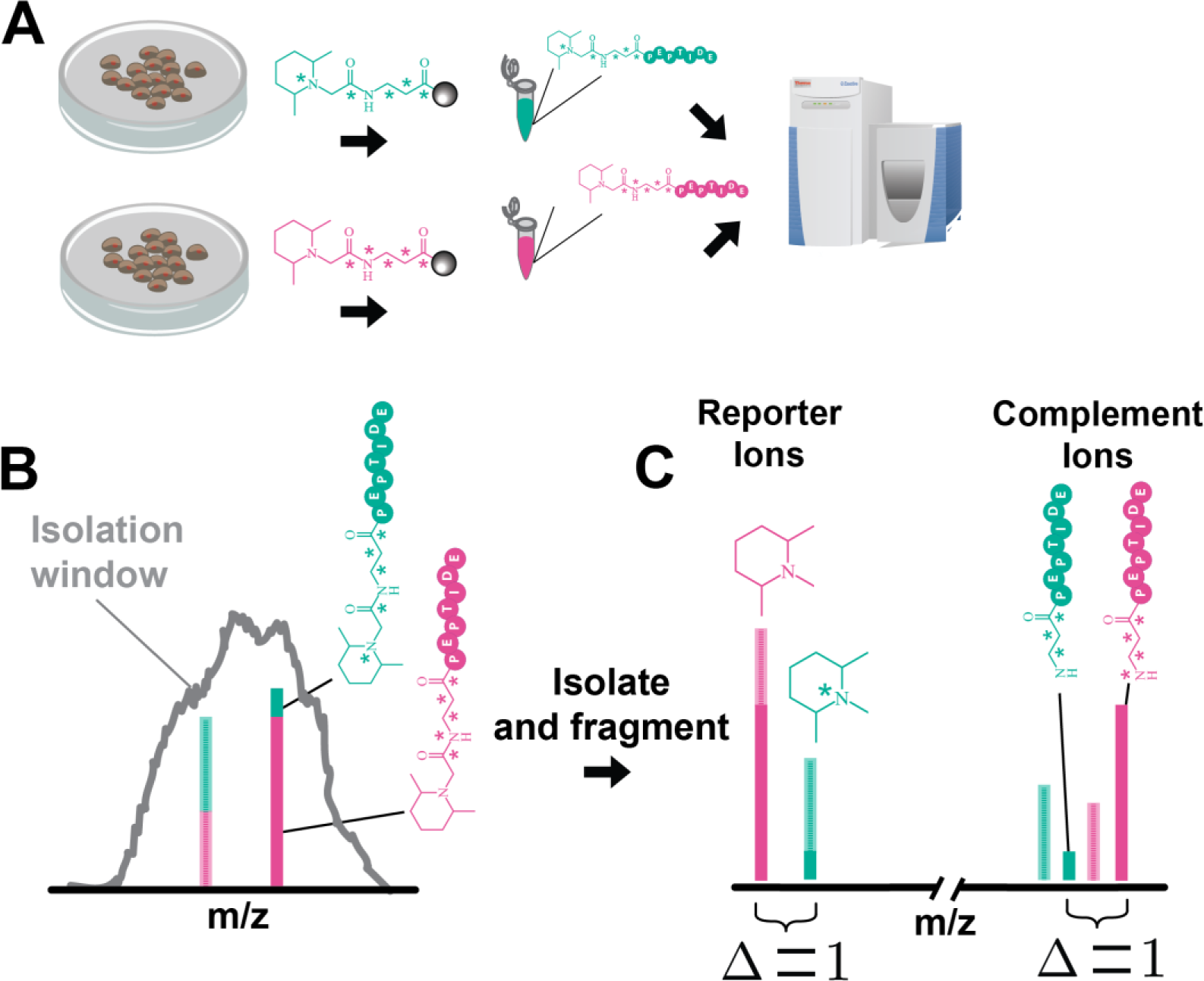
Concept of quantification with the complement reporter ions. **A)** Proteins from complex mixtures are digested into peptides and labeled with isobaric tags (green or pink) that contain heavy isotopes (stars). These tags serve as barcodes for different conditions. After tagging different conditions (replicates, time points etc.) are mixed together and ionized onto a mass spectrometer. **B)** In the MS1 scan different isobaric tags have the same mass, thus peptide peaks from different conditions are indistinguishable. A target peptide (dark green/pink peak) is isolated, but typically at least one other peptide (shorter, opaque green/pink peak) is also simultaneously co-isolated. **C)** The isolated peptides are fragmented, and in the case of TMT-MS2, the signal from the reporter ions in each condition is distinguishable by mass and can be quantified. However, since an additional peptide was co-isolated, the signal from each peptide sum producing a distorted ratio. Alternatively, with TMTc, the remaining “complement ion” portion of the isobaric tag attached to the peptide is quantified. The peptide itself confers a unique mass to each peptide after fragmentation. Co-isolated peptides will have a slightly different mass that is typically distinguishable, enabling accurate quantification even when other peptides are co-isolated and fragmented.

Measurements with this simple implementation of multiplexed proteomics (TMT-MS2) are typically severely distorted.^6^ In complex mixtures the MS2 spectrum typically contains a mixture of fragments from the peptide of interest and other co-isolated peptides (Fig. 1C). Therefore, the measured signal is actually a combination of reporter ions from the identified peptide and reporter ions from other peptides. Here-in we will refer to ratio distortion that arises from reporter ions of co-isolated peptides as interference. Interference can be minimized by an additional isolation and fragmentation of b- and y-ions in the MS3 scan.^6b, 7^ This approach, termed MultiNotch MS3 (TMT-MS3), has been commercialized on the Orbitrap Fusion and Lumos as “synchronous precursor selection-based MS3” and allows the routine quantification of ~8,000 proteins. The improvements from TMT-MS3 are now considered the current state-of-the-art for multiplexed proteomics, and have allowed detection of protein abundance changes of 10% as highly significant.^8^

While the MS3 approach is a significant advance, it does not completely eliminate interference; especially for low abundant peptides where interference remains a major problem.^8^

Furthermore, additional MS scans result in a slow duty cycle and loss of ions, which manifest in loss of sensitivity. Lastly, TMT-MS3 data can only be obtained on expensive and complex instrumentation, which are comparatively difficult to maintain. Despite the clear progress TMT- MS3 introduced, TMT-MS2 is still used an estimated ~5 times more often in academic studies, based on the number of times each method was cited in 2016.^8,9^

We have previously devised a system of protein quantification using the complement reporter ions (TMTc).^8, 10^ This method nearly eliminates interference. When TMT-labeled peptides fragment at the MS2 stage of mass analysis to produce the low m/z “reporter ions” discussed above, additional complement reporter ions are formed as a result of the intact peptide remaining fused to the mass balancing region of the TMT-tag (Fig 1C). The TMTc ions encode different experimental conditions in the same way the low m/z reporter ions do, with the added benefit that the TMTc ion’s mass is different for each peptide. Therefore, accurate quantification is possible even if other peptides are co-isolated into the same MS2 spectrum.

TMTc is better in distinguishing true signal from interference because the ability to distinguish peaks in mass analyzers in the MS2 scan is typically ~100-fold higher than the narrowest possible isolation window that can be used for isolating ions into the MS2 scan. Because TMTc quantification does not need the additional gas-phase isolation-step of the slow MS3 scan, it holds potential to be significantly more sensitive and is compatible with comparatively simple instruments like iontrap Orbitrap, quadrupole Orbitraps, and Q-TOFs.

Despite these obvious advantages, published TMTc methodology is limited by comparatively low measurement precision and inefficient TMTc ion formation, which reduces sensitivity. While TMTc is already able to provide accurate data that has comparable sensitivity as data gathered using TMT-MS3,^8^ specific improvements are needed to fully exploit its potential. Here, we introduce these advancements in a method we term TMTc+. We modified sample preparation and instruments methods to favor the complement ion formation. Further, we included the shape of the isolation window into the deconvolution algorithm, which drastically increases precision. At the same time the improved method maintains the superb ability to distinguish signal from chemical noise. Thus, TMTc+ enables accurate, sensitive, and precise multiplexed proteomics that is compatible with widely distributed instrumentation.

## Results and Discussion

### Improving the precision of the complement reporter ion approach: TMTc+

One major obstacle for TMTc in its published form is poor measurement precision: there is a large unbiased spread in measurements that surround the true ratio. The main source of imprecision in the TMTc method arises during the deconvolution procedure. Consider a TMTc experiment where equal amounts of a single peptide in five different conditions are each labeled with a different TMT tag and analyzed (Fig S1A). Since 1% of all carbon is ^13^C, a single peptide exists as an isotopic envelope, a distribution with several different peaks each differing by one Dalton. Within each peak in the isotopic envelope, the relative abundance of each condition for the peptide is equal, however in reality these ratios are unknown. In the limiting case where only a single peak, e.g. the pseudo-monoisotopic peak (M), is isolated and fragmented (Fig S1B), the correct ratios can be read out directly (after minor correction for isotopic impurities in the TMT tags). If the isolation is pure, the correct ratios can also be determined from another peak within the envelope (e.g. M+1 peak), however note the mass offset of each condition is shifted each by 1 Da because the peak selected from the precursor envelope was the M+1 peak (Fig S1C). In the case where both the M+0 and M+1 peak are simultaneously isolated and fragmented the complement ions from each condition offset, and a convolution occurs: the observed ratios are now incorrect (Fig S1D). The true underlying ratios can be estimated using the theoretical distribution of peak heights in the precursor envelope, but this process comes with loss of precision.^10^ In its published form, TMTc isolates and fragments the entire precursor envelope (Fig S1E).

We hypothesized that using a small isolation window and including its shape in the deconvolution algorithm would lead to improved measurement precision due to the simpler deconvolution process. We measured the shape of the isolation windows for various settings (e.g. 0.5 Th and 1.0 Th) by measuring the obtained signal of infused MRFA peptide, while we scanned the radio-frequency of the quadrupole around to shift the isolation window around the peptide’s mass. We show a subset of these measured windows in (Fig S2) and provide a description of windows from 0.4 Th to 2.0 Th in 0.1 Th increments in Table S1. Knowing the shape of the isolation window for a given width used by the quadrupole during isolation allows us to model the isolation step in the deconvolution algorithm by how much of each peak was isolated from the precursor envelope. We term this approach TMTc+.

To evaluate the measurement precision of TMTc and TMTc+ experimentally, we labeled equal amounts of peptides from HeLa lysates with five different TMT-tags (Fig. 2A). The expected true ratios for this sample are 1:1:1:1:1. When we analyze the sample with the published TMTc method that isolates the entire precursor envelope, we obtain a fairly wide spread around the true answer with a mean coefficient of variation (CV) of 24% (Fig. 2B). In contrast, when using TMTc+ with a much smaller 0.5 Th isolation window (Fig 2C), typically a single peak can be purely isolated, and the mean CV improves drastically to 8%. Importantly, the TMTc+ method is generalizable and can accommodate isolation windows of arbitrary shapes. Thus, larger windows e.g. 1.0 Th can also be used and significant improvements in the mean CV relative to the published TMTc method are still observed (Fig 2D). We expect a trade-off between measurement precision and sensitivity, with narrower isolation windows increasing the precision of each measurement but coming at the expense of less signal. All of the following studies in this paper except those in figure 5 use a 0.4 Th isolation window, the narrowest window currently accessible. This was chosen to test TMTc+ under the most challenging conditions possible: conditions that in principle give the best measurement quality but are the least sensitive.

**Figure 2:**
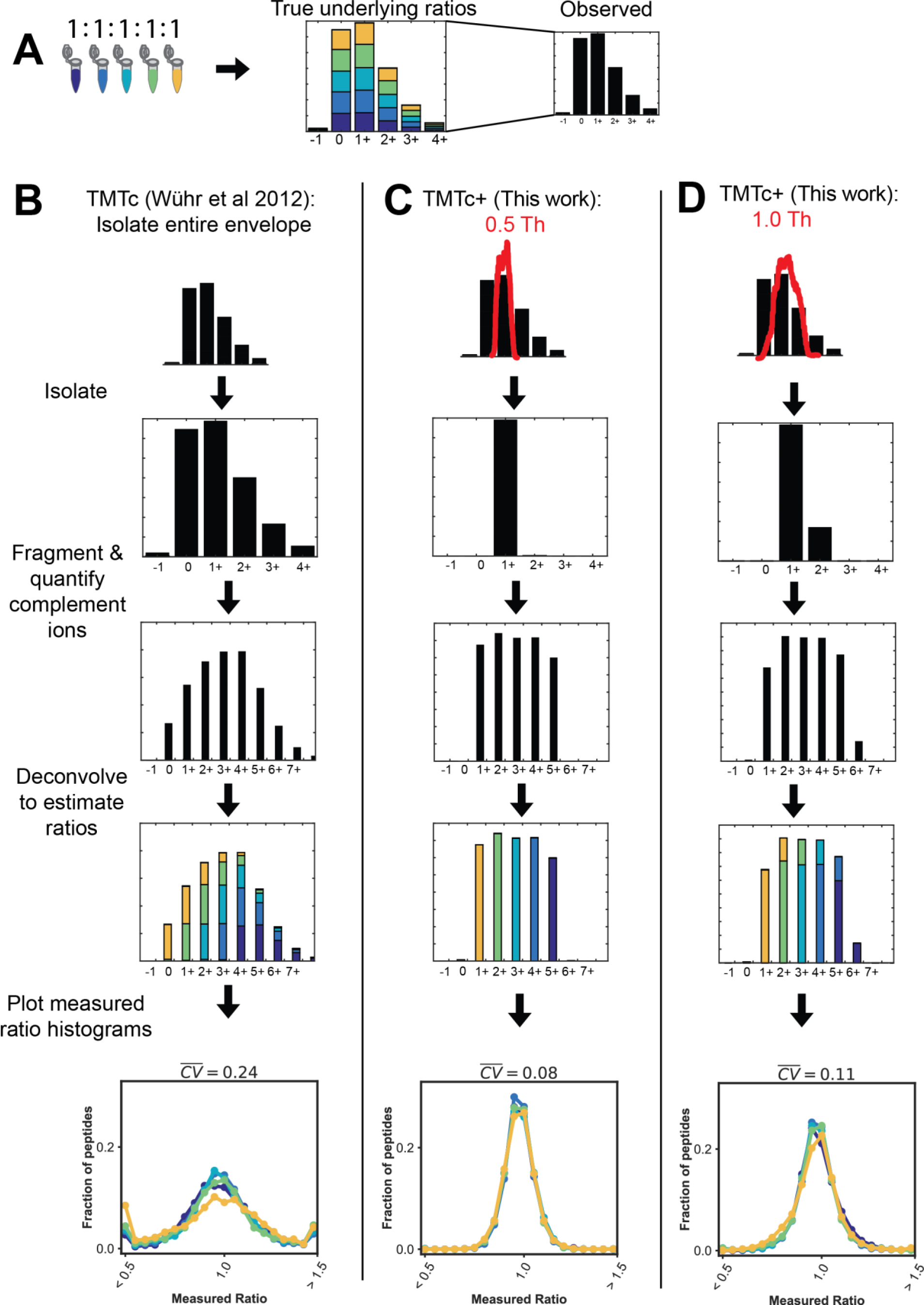
Modeling the isolation step in the deconvolution algorithm increases measurement precision. **A)** Five different TMT reagents are used to barcode five identical samples of peptides from a HeLa lysate. An example of the isotopic envelope of an intact peptide (precursor) is shown. The true underlying ratios are shown in color, but the mass spectrometer is blind to the barcoding and only what is shown in black is observed. Note that in the MS1 before any fragmentation has occurred that the true ratio of 1:1:1:1:1 is present in each peak of the envelope. **B)** In the published form of TMTc the entire monoisotopic envelope is first isolated and then fragmented. The peaks corresponding to the complement reporter ions are identified and the relative abundances are determined. The simultaneous fragmentation of multiple peaks from the precursor convolves the data. Intuition for this convolvement is provided in Fig S1. Using the known theoretical distribution of charge states from the precursor, the amount of signal in each mass offset that belongs to each condition can be estimated using a least-squares optimization.^10^ This process is done for thousands of peptides and the resulting histograms of the measured ratio are shown, resulting in a mean CV of 24%. **C)** With TMTc+ the shape of a much narrower (e.g. 0.5 Th) isolation window (red) is measured and used. The shape and position of this window is incorporated into the deconvolution algorithm. With narrower isolation windows deconvolution becomes easier and precision is gained. In the extreme case where only a single peak from the envelope is isolated, the TMTc+ algorithm has to only calculate away isotopic impurities from the TMT-tag. The median CV on the peptide level improves to 8%. **D)** TMTc+ can accommodate isolation windows of any size as long as their shape has been measured. In the case where a 1.0 Th window is used (red), a small amount of the M+2 peak is isolated in addition to the M+1 peak. Using the shape of this moderately sized isolation window, the ratios can be deconvolved and a significant improvement in precision (mean CV of 11%) is still observed relative to when the entire isotopic envelope is isolated with previously published TMTc.

### Comparison of measurement accuracy between TMTc+, TMT-MS2, and TMT-MS3

In the previous section we showed the improvements in measurement precision the TMTc+ algorithm is able to generate. Here we evaluate the ability of TMTc+ to distinguish real signal from background in a complex sample where many peptides can be co-isolated along with the peptide of interest. These co-isolated peptides cause a significant amount of interference for TMT-MS2, and were the motivation for the development of MultiNotch MS3 (TMT-MS3), which mitigates this problem and is the current state of the art.^6b, 7^ To assess the ability to distinguish true signal from background we developed a two-species standard (Fig 3A). The two species standard consists of human peptides from HeLa cells labeled with five different TMT-tags and mixed at a ratio of 1:1:1:1:1. We add yeast peptides at 10x lower levels with a mixing ratio of 1:0:1:0:1. The 10x difference in abundance levels allows us to approximate that the majority of interference is contributed by HeLa peptides. To evaluate the ability to measure signal relative to chemical noise (CSN), we measure the relative signal for unique yeast peptides in a condition containing yeast and human peptides (TMT 126) relative to a condition with only human peptides (TMT 127) (Fig 3A). A perfect measurement would yield a CSN of infinity. Importantly, the TMT 126 and TMT 127 conditions used in this assay represent the most difficult scenario for TMTc+. These channels only differ by 1 Da and are thus the conditions “right next to each other” in the complement reporter region of the mass spectrum (Fig S1). Thus, any signal that is not correctly deconvolved will “spill over” and decrease the CSN of TMTc+ (e.g. see Fig. S1D).

**Figure 3:**
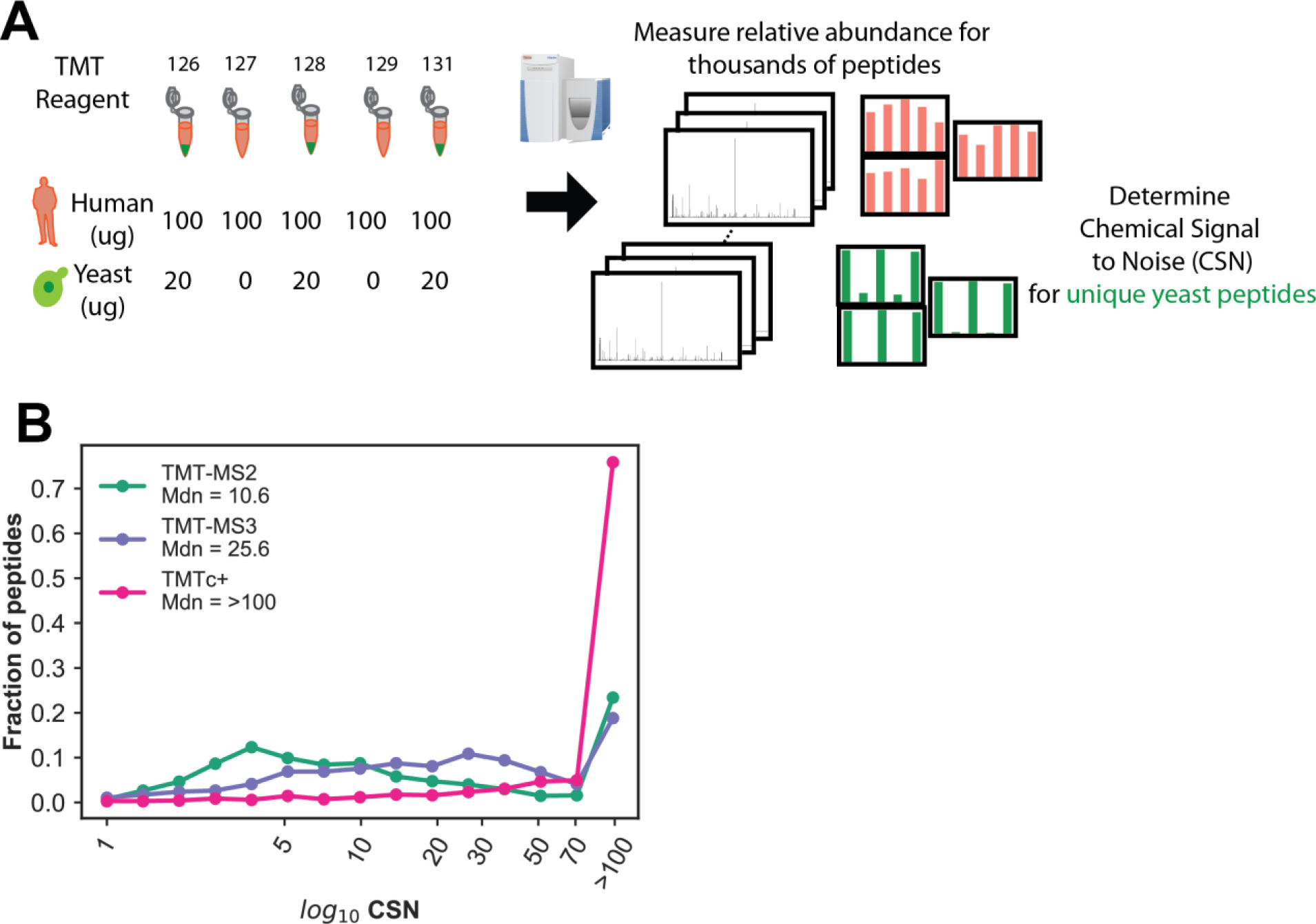
Signal to noise comparison between TMTc+, TMT-MS2, and TMT-MS3. **A)** A yeast / human standard was designed to assay the chemical signal to noise of TMTc+ relative to other multiplexed proteomics methods in use. Yeast (green) is only labeled with 3 of the 5 TMT reagents used, whereas human (pink) is labeled with all 5. The ratio of 126 TMT / 127 TMT for unique yeast peptides is used to assay the chemical signal to noise (CSN). A perfect ^measurement shows an infinite change. **B**) log10 CSN histogram of unique yeast peptides using^ TMT-MS2, TMT-MS3, and TMTc+ from a 90 minute reverse phase fractionated sample. Bins are indicated with a circle. Due to restraints on ion counting statistics, any CSN larger than 100 was plotted as 100.

Figure 3B shows the CSN distributions for TMTc+ relative to TMT-MS3 and TMT-MS2. Consistent with previous publications,^6b, 7^ the median CSN of TMT-MS3 is 25.6 and superior to that of TMT-MS2, which has a median CSN of 10.6. In contrast to both of these methods, we observed more than 70% of all peptide measurements made with TMTc+ have CSNs that are >100. The ability of TMTc+ to remove interference is superior to TMT-MS2, and despite the omission of an additional gas-phase purification step, even better than TMT-MS3. We observe similar results for the CSN of TMTc+ if we use the 128 and 129 channels instead of 126 and 127 (Fig S3). When comparing the precision of these three methods, we note that TMTc+ has slightly lower but comparable precision (Fig S4).

### Comparison of measurement sensitivity between TMTc+, TMT-MS2, and TMT-MS3

Another disadvantage of the published TMTc method was lack of sensitivity, mostly due to inefficient formation of complement reporter ions. This was most pronounced for longer peptides with higher charge states.^11^ To address this problem we modified sample preparation by switching from a LysC only digest to Trypsin/LysC, which produces shorter peptides with fewer missed cleavages. Additionally, we added 2% DMSO to the ionization medium to coalesce charge states towards 2+, and optimized the instrument method.^12^ For a detailed sample preparation protocol and method setup please refer to the materials and methods section. To evaluate and compare how many proteins we could quantify with different multiplexed methods, we analyzed a five-plex HeLa lysate (containing only human and no yeast proteins) with TMTc+, TMT-MS2, and TMT-MS3. For this comparison we used a stringent 1% FDR on the protein level,^13^ and count only the minimal number of proteins required to explain all observed peptides.^14^ TMTc+ is able to measure ~700 more proteins for 24 90 minute analyses of pre-fractionted samples relative to TMT-MS3. TMTc+ is likely able to quantify more proteins than TMT-MS3 due to the faster duty cycle and the omission of the extra gas-phase purification step (Fig. 4A). The number of quantified proteins with TMTc+ even slightly surpasses the semi-quantitative TMT-MS2 method (Fig. 4A). The advantages/disadvantages of the methods seem to roughly cancel each other: the commercial TMT-tag was designed to favor the production of low m/z reporter ions instead of complement reporter ions during fragmentation. Consequently, TMTc+ methods must use longer ion-injection times for each peptide to obtain the same number of ions. This lowers the duty cycle of TMTc+, and a larger fraction of spectra cannot be used for quantification. On the other hand, many spectra for TMT-MS2 (and TMT-MS3) are filtered out due to obvious co-isolation of other peptides within the MS1 isolation window (as previously described we require an isolation specificity of 75%). For TMTc+, we don’t need to use this filtering step, as we are typically able to distinguish signal from the peptide of interest from co-isolated peptides in the MS2 spectrum (Fig. 3). Thus, even when the narrowest isolation window of 0.4 Th is used with TMTc+, the method is as or more sensitive than existing methods. For all three methods, one can reduce the number of reverse phase fractions analyzed to obtain a proteome at a reduced sensitivity in exchange for saving instrument time (Fig 4B.)

**Figure 4:**
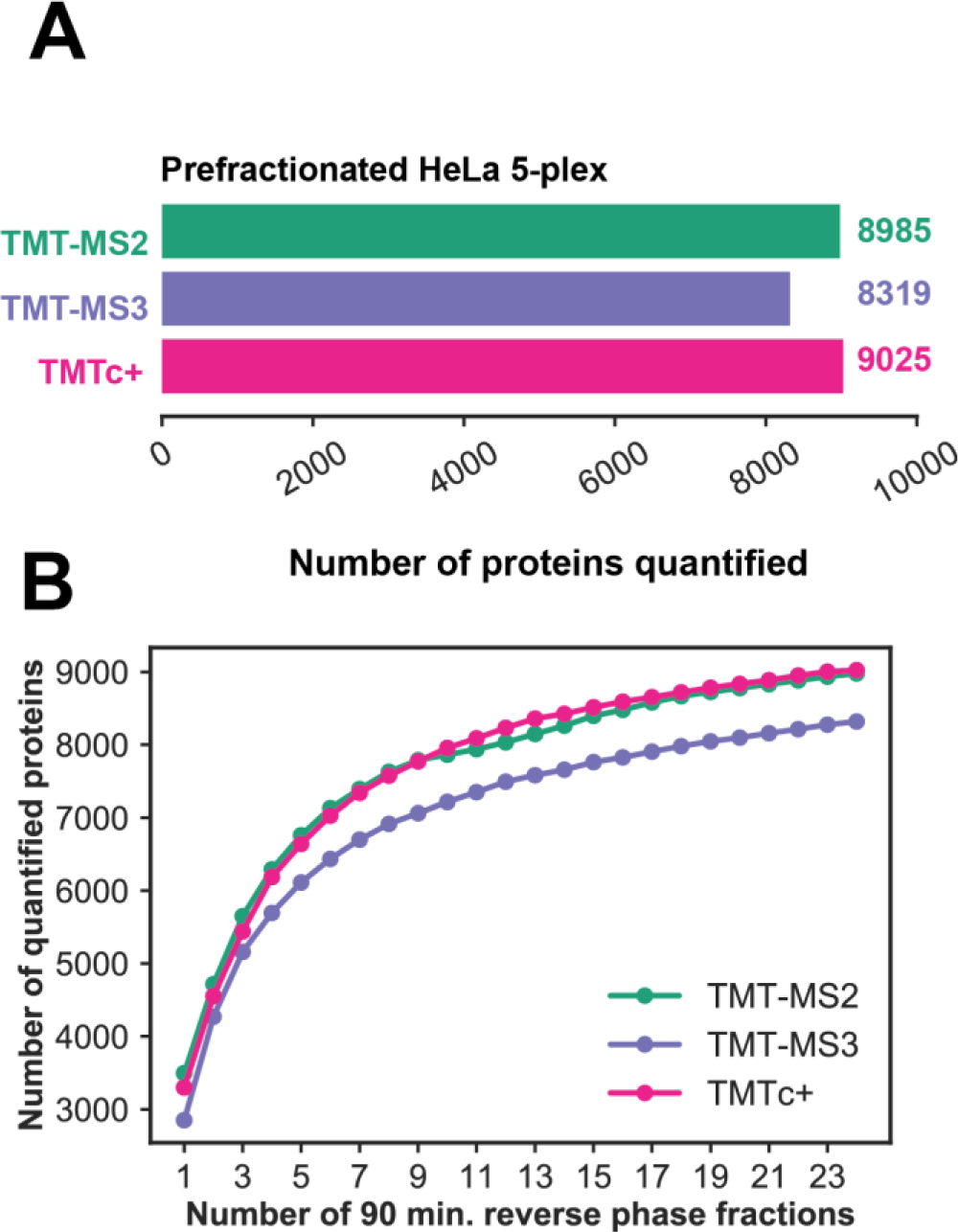
Comparison of Sensitivity for different multiplexed proteomics methods. **A)** Number of proteins quantified at a 1% FDR on the protein level after analyzing 24 90 minute reverse phase fractionated samples of a 1:1:1:1:1 TMT tagged HeLa standard. TMT-MS2 and MultiNotch-MS3 measurements were filtered to an isolation specificity > 75% as previously described.^7–8, 18^ However, this was not necessary for TMTc+. **B)** Number of proteins quantified at a 1% FDR on the protein level as a function of the number of 90 minute reverse phase fractionated samples analyzed with each method.

### Improving the robustness and accessibility of TMTc+

So far, we have demonstrated TMTc+ can produce superb measurement quality combined with very high sensitivity. Additionally, one of the most attractive points of TMTc+ is that it does not require an ion-trap and is compatible with quadrupole Orbitrap instruments, which are at least 10x more prevalent in the field than TMT-MS3 compatible instruments based on the number of datasets that have been deposited to the proteomics database Proteome Xchange. TMTc+ can be used by many labs which currently only have access to instrumentation that can perform the TMT-MS2 method. However, an important requirement for TMTc+ is that we use the shape of a highly reproducible isolation window into the deconvolution algorithm. It is possible that due to poor calibration or older instrumentation, that this isolation window is not well defined or changes drastically as function of m/z (Fig S2). To assess the dependency of TMTc+ on the correct isolation window used we acquired data with a 0.7 Th isolation window, but performed the subsequent deconvolution assuming a 0.4 Th isolation window was used. As expected data quality suffers (Fig 5A). Upon inspection of individual MS2 spectra, we observed that the vast majority of spectra contained unfragmented precursor ions, which we term surviving precursor (Fig. 5B). We hypothesized that the ratio of each peak in the surviving precursor can be used as a direct readout for how much of each peak from the precursor envelope was isolated and fragmented. When performing the deconvolution using the surviving precursor ratios of the same 0.7 Th data acquired as in (Fig. 5A) the median CSN improved from 14.6 to 28.9 (Fig 5C). The surviving precursor can be used for deconvolution of almost every spectra (Fig 5D).

**Figure 5:**
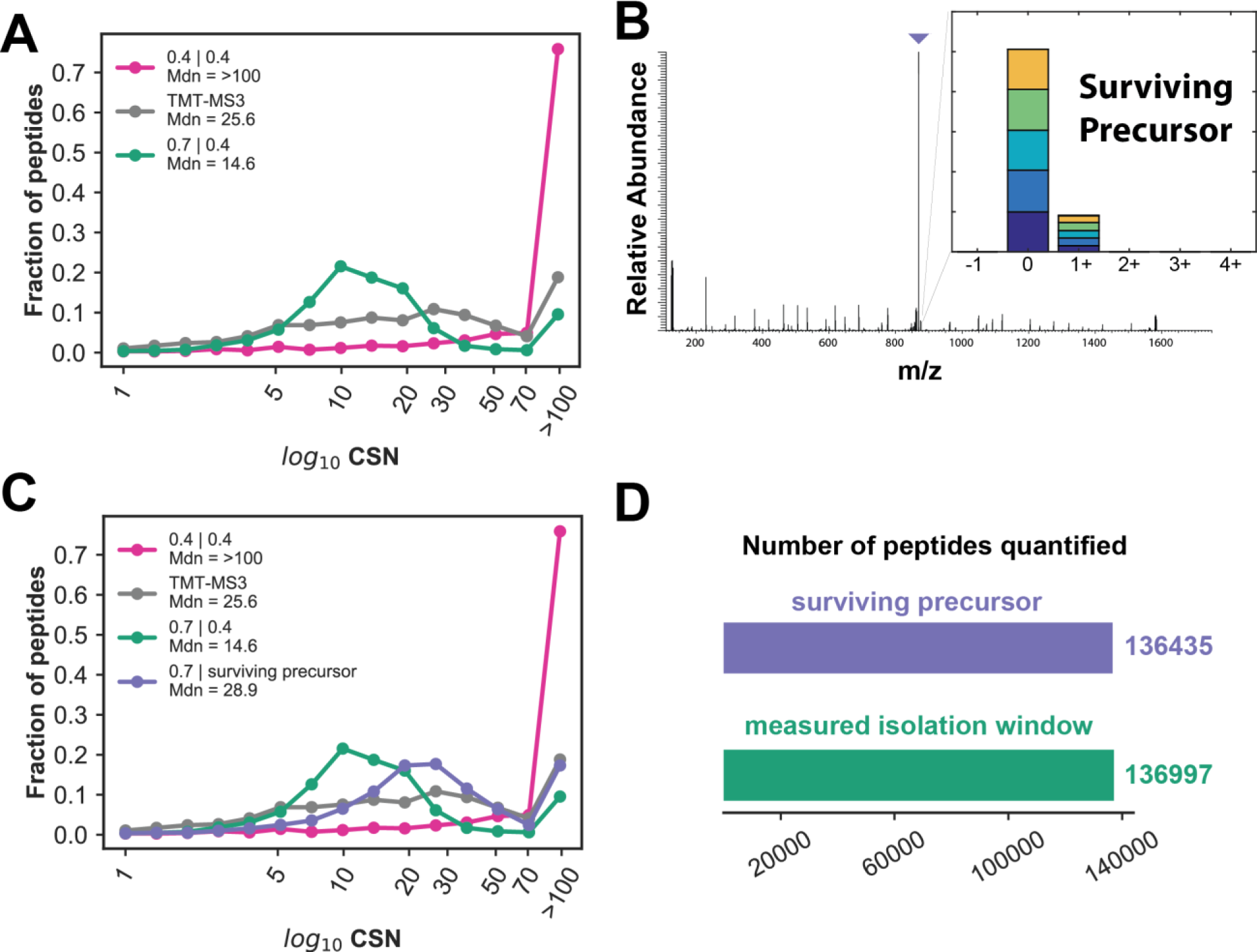
Incorporation of the surviving precursor monoisotopic envelope shape into the deconvolution algorithm increases robustness of TMTc+. **A)** Histograms of CSN of TMTc+ where the human / yeast standard was analyzed using different isolation windows on the instrument that are deconvolved using either proper or improper isolation window (IW) shapes (IW used on instrument | IW used during deconvolution). In the best case scenario for TMTc+, the narrowest isolation window (0.4 Th) is used and the shape of the 0.4 m/z isolation window is used during the deconvolution (pink, (0.4 Th | 0.4 Th)). To mimic a poorly calibrated instrument a 0.7 Th isolation window was used by the instrument, but the data was deconvolved assuming the shape of the isolation window was 0.4 Th (green (0.7 Th | 0.4 Th)). Shown for reference is the CSN for TMT-MS3 (grey). **B)** Representative MS2 spectra with a TMTc+ method. The tallest peak (purple arrow) is the "surviving precursor", intact precursor from the MS1 that was not fragmented. Close inspection of the surviving precursor reveals that the ratios of each peak isolated from the pseudo-monoisotopic envelope in the MS1 (inset, colored bars) can be quantified. **C)** Same data as in A) but the same sample that was analyzed on the instrument with a 0.7 Th IW is now deconvolved using the ratios from the surviving precursor in the MS2 of each peptide instead of incorrectly assuming the isolation window used was the shape of 0.4 Th (purple (0.7 Th | surviving precursor)). An increase in CSN is observed. **D)** Number of peptides that are quantified from 24 90 minute reverse phase fractions using TMTc+ with either the surviving precursor approach (purple) or with the measured isolation window (green).

Despite this progress, we were not able to reach the same quality of data as what is observed when the correct theoretical isolation window is used. We believe this is due to the introduction of noise with the measurement of the surviving precursor. In summary, we believe that the best possible data can be acquired by using the theoretical isolation window of a well calibrated instrument. Nevertheless, using the surviving precursor might be an attractive alternative which can increase robustness and still provide measurements that are equivalent to the current state of the art. We expect this method might be particularly useful for iontrap-Orbitrap instruments (e.g. Orbitrap Velos) where space charging effects in the linear ion trap lead to poor reproducibility of the centering of the isolation window.

### Comparison of TMTc+ with label free quantification

Despite the theoretical advantages of multiplexing methods, practical difficulties such as instrument compatibility and interference have hindered adoption. The most commonly used form of quantitative proteomics is “label free”, where multiple unlabeled samples are analyzed in succession on separate runs of a mass spectrometer, and then the intensities of their peaks from each run are compared.^3^ We believe these former advantages of label free approaches relative to multiplexed ones are diminished by TMTc+. We assayed the sensitivity and precision of both methods by analyzing five identical samples of the same HeLa cell lysate (Fig 6A).

**Figure 6:**
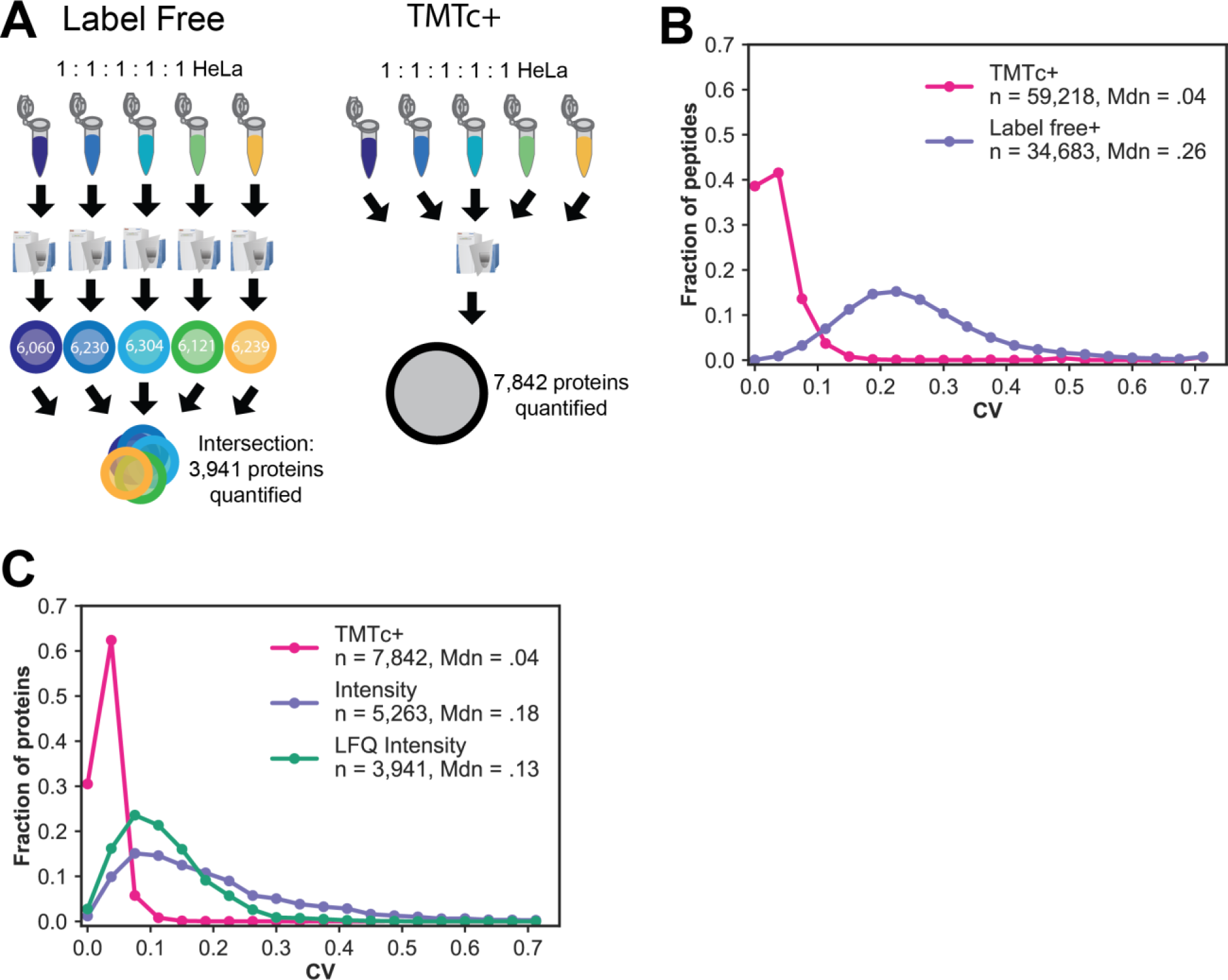
Comparison of TMTc+ with label free proteomics. **A)** A sample of digested HeLa peptides was either labeled with five different TMT reagents and combined to be analyzed by TMTc+, or analyzed in succession via one-shot label free proteomics^3^ for the same amount of instrument time (15 hours total). An additional 3,901 proteins are quantified in all five samples using TMTc+. **B)** Distribution of coefficients of variation at the peptide level with each method. TMTc+ (pink) quantifies an additional 24,535 peptides relative to label free (purple) and has a median CV of 4% relative to a median CV of 26% with label free quantification. **C)** Peptide measurements were aggregated to the protein level (1% FDR) either by summing counts of ions in each sample for TMTc+ (pink), or by two different established methods for label free (green and purple).^3^ TMTc+ quantifies thousands of additional proteins compared to both quantitative label free approaches, and has a lower median CV (4%) relative to the label free approaches (13% and 18% respectively).

Using the same amount of instrument time for both methods, we identified a total of 5,263 proteins at a 1% FDR in all five label free runs. Of these, 5,263 were quantified using the intensity metric in MaxQuant, whereas 3,941 were quantified using the LFQ quantification.^3^ In the same amount of instrument time, TMTc+ quantified 7,842 proteins at a 1% FDR. The substantial increase of proteins quantified across all five conditions with TMTc+ relative to label free can be attributed to 1) using the same total amount of instrument time, the co-analysis of five samples simultaneously with TMTc+ at once allows the analysis of multiple pre-fractionated peptide samples. 2) The multiplexed nature of TMTc+ results in peptides, and thus proteins, being simultaneously identified in all five conditions. With TMTc+ there are no missing values to remove. Last, the precision of each method was compared using the same HeLa samples on both the peptide level (Fig 6B) as well as the protein level (Fig 6C). On the peptide level the median CV for TMTC+ is 4% whereas for label free it is 26%. When peptide measurements are aggregated to the protein level the median CV for TMTc+ is 4%, with essentially no proteins having a CV higher than 10% for 7,842 proteins. In contrast, with the label free Intensity quantitative metric 5,263 proteins can be quantified with a median CV of 18%. The label free CV can be improved to 13% using the LFQ Intensity metric,^3^ but the precision is still comparatively poor and ~4,000 less proteins are quantified.

Taken together, we believe that TMTc+ provides an attractive alternative to label-free quantification. It results in superior measurement precision, higher sensitivity, and removes the interference, which is a major problem with TMT-MS2. A major hindrance for the adaption of accurate multiplexed proteomics has been the incompatibility of TMT-MS3 with the most widely used instrumentation. We hope that TMTc+ will help to overcome this limitation.

While TMTc+ is now at least as sensitive as other multiplexing methods in use, we believe that there is promise for even larger gains in sensitivity to be achieved. The current TMT tags used for the method were not designed with TMTc+ in mind and form the complement reporter ion inefficiently. Development of new TMT reagents with more selective fragmentation that favor the complement reporter fragment might yield up to an additional order of magnitude in signal for the method. We believe that the inefficient ion formation is the main reason for the slightly poorer measurement precision observed for TMTc+ compared to TMT-MS2 and TMT-MS3 (Fig S4). We expect a more suitable tag to further improve TMTc+’s measurement precision. We noticed the very efficient formation of complement reporter ions in an isobaric tag that was designed for a different purpose.^15^ Currently we only attempt to identify and quantify peptides with a 2+ charge state because the spacing between peptides with a 3+ charge state is too narrow to use a 0.4 Th isolation window, the narrowest isolation window currently available on quadrupoles from Thermo. We suspect that if it were possible to use narrower isolation windows, e.g. 0.2 Th, it would be possible to also quantify most 3+ charge state peptides, while maintaining superb measurement precision. Quantification of 3+ peptides (and even higher charge states) would of special interest for certain endogenous post-translational modifications as well as derivatization with different chemical compounds. Finally, due to the unique m/z of each complement reporter fragment due to the peptide, it is possible to quantify multiple peptides in the same spectra and therefore paralyze data acquisition.^10^

Compared to current approaches, the remaining limitation with TMTc+ is its multiplexing capacity. With the commercially available TMT reagents, the maximum amount of different encoded conditions is five. This compares unfavorably to the 11 channels that are currently accessible with TMT-MS3, which also produces excellent data. The reduced number of channels is due to the positioning of the heavy isotopes relative to the two breakage points in the TMT-tag, and the lower mass resolution obtainable at higher m/z. Currently, we are unable to distinguish the neutron masses of nitrogen or carbon in complement reporter ions. Nevertheless, using the current approach, a new set of reagents encoding ~ten different conditions each with one Dalton spacing seems feasible. We hope that the promising results shown in this paper will motivate the generation of such tags. The multiplexing number could further increase with super-resolution approaches that would, in principle, allow the use of neutron encoding of conditions for complement reporter ions.^16^

Lastly, the unique ability of TMTc+ to distinguish signal from noise might prove particularly useful for targeted multiplexed proteomics.^17^

## Conclusion

Our recent improvements to the complement reporter approach result in a method we term TMTc+. This method produces measurements with significantly better signal to noise than the current state of the art, TMT-MS3. TMTc+ is compatible with at least 10 times more mass spectrometers in use relative to TMT-MS3 at the time of this publication. An important aspect of TMTc+ is that our approach can be generalized to use isolation windows of any shape. This might be especially important for older instruments that have lower sensitivity. The deconvolution software used in this paper is freely available upon request for all academic users. The software can be used as an add-on to existing excellent mass spectrometry analysis pipelines such as MaxQuant or Proteome Discoverer.

## Acknowledgement

We would like to thank Graeme McAlister (Thermo Fisher) for comments and suggestions and the software to measure the isolation window. We would like to thank Ramin Rad for converting the TMTc+ code into a standalone software package. M.S. was supported by Ruth L Kirschstein National Institutes of Health F31 Predoctoral Fellowship 5F31GM116451. This work was supported by Princeton startup funding and the LSI collaboration fund.

